# Double trouble or a blessing in disguise? Co-infection of potato with the causal agents of late and early blight

**DOI:** 10.1101/2022.05.31.494103

**Authors:** S.M. Brouwer, P.J. Wolters, E. Andreasson, E. Liljeroth, V.G.A.A. Vleeshouwers, L.J. Grenville-Briggs

## Abstract

The simultaneous occurrence of multiple diseases is an understudied area in plant pathology; however, studies of animal and human diseases have shown that the presence of multiple pathogens can impact virulence, and the course of disease development. Furthermore, they also present an important driver of epidemiological dynamics. Global potato production is plagued by multiple pathogens, amongst which are *Phytophthora infestans* and *Alternaria solani*, the causal agents of potato late and early blight respectively. Both these pathogens have different lifestyles and are successful pathogens of potato, but despite observations of both pathogens infecting potato simultaneously in field conditions, the tripartite interactions between potato and these two pathogens are so far, poorly understood. Here we studied the interaction of *A. solani* and *P. infestans* first *in vitro* and subsequently *in planta* both in laboratory and field settings. We found that *A. solani* can inhibit *P. infestans* both in terms of growth *in vitro* and infection of potato, both in laboratory experiments and in an agriculturally relevant field setting.

## Introduction

Worldwide, the production of potato, *Solanum tuberosum* L., is affected by plant pathogens causing diseases. The notorious late blight pathogen *Phytophthora infestans* (Mont.) de Bary, that infamously destroyed potato production in many countries in the 19^th^ century and launched the study of phytopathology, is still the number one potato pathogen (Judelson & Blanco, 2005, Savary et al., 2019). Early blight, caused by the fungus *Alternaria solani* Soraurer, has the potential to reduce the yields by half if left uncontrolled (Leiminger and Hausladen, 2012). Currently, both potato diseases are controlled mainly by repeated applications of synthetic fungicides. Yet, fungicide resistance, new regulations concerning fungicide use, and a changing climate are all factors that challenge current disease control strategies (Elad and Pertot, 2014, Landschoot et al., 2017, Schepers et al., 2018, Odilbekov et al., 2019).

Plant pathogens such as *Phytophthora infestans* and *Alternaria solani*, are devastating and notorious pathogens plaguing potato production individually (Savary et al., 2019), however, in an agricultural field multiple pathogens are present or can arrive at any time. Yet, the effect and interactions of unrelated fungal and oomycete pathogen species infecting the same host plants is largely unknown and understudied. Nonetheless, the infection of multiple unrelated pathogen species has been shown to result in strong effects in animal and human pathology (Laminichhane &Venturi, 2015, Tollenaere *et al*., 2016). The challenge that multiple pathogens can infect a host plant or host population, has been identified as one of the thirteen challenges in modelling plant diseases (Cunniffe *et al*., 2015). Additionally, it has been suggested that co-infected plants should be targeted in disease control programs since co-infection can be an important driver of epidemiological dynamics (Susi *et al*., 2015).

*A. solani* is classified as a necrotrophic pathogen but, according to previous studies, forms appressoria (Dita *et al*., 2007). The formation of an appressorium to infect the plant without causing considerable damage to the plant tissue is a strategy commonly used by biotrophic and hemibiotrophic pathogens but also employed by necrotrophic pathogens (Mendgen & Hahn, 2002, Prins et al., 2000). A study using hyperspectral spectroscopy collecting reflectance data between 400 and 2400 nm for the detection of both *P. infestans* and *A. solani* in potato leaves, was able to detect distinct changes in the leaf reflectance spectra for both *A. solani* and *P. infestans* before lesions appeared (Gold et al., 2020), indicating that *A. solani* and *P. infestans* trigger different responses in the plant during early infection before the development of characteristic early or late blight specific lesions. *P. infestans* is a hemibiotrophic pathogen that in a susceptible interaction suppresses plant defences by the use of effector proteins, allowing biotrophic growth, however, at a later stage, the switch to necrotrophy is made and necrosis inducing effectors, and other molecules, are excreted (Lee & Rose, 2010).

When multiple pathogens require similar resources from the plant, this could influence co-infection/ cohabitation dynamics. For example, if one of the pathogens triggers a response in the plant, this could be beneficial or negative for infection by the other pathogen (Tollenaere *et al*., 2016). When *Arabidopsis thaliana* is inoculated with *Pseudomonas syringae* and subsequently challenged with *Alternaria brassicicola* on the same leaf, larger *Alternaria* lesions were observed than for the control. However, inoculation of *P. syringae* and *A. brassicicola* on different leaves did not lead to significant differences in lesion size (Spoel *et al*., 2007) even though competition between pathogens with a similar lifestyle might be expected. *P. infestans* is not usually able to infect *A. thaliana*, which is therefore classified as a nonhost for this pathogen. However, Belhaj et al. (2017) showed that when *A. thaliana* was first infected and colonised by the oomycete *Albugo laibachii* this rendered the plants susceptible to subsequent *P. infestans* infection. Moreover, both pathogens were observed to form haustoria in the same cells, indicating that the formation of a feeding structure by one pathogen did not limit formation of a feeding structure by the second pathogen. Furthermore, the *A. laibachii* infected *A. thaliana* plants were not found to be more susceptible to *Blumeria graminis* f.sp. *hordei* and *Phakopsora pachyrhizi*, two other non - *A. thaliana* adapted pathogens. The effect of multiple pathogens present in a host plant at the same time, is thus dependent on the specific combination of pathogens. Studies on these compatible or incompatible tripartite or multiparty interactions between plants and pathogens help to further elucidate susceptibility and resistance mechanisms, offering important insights into how disease dynamics may play out in an agricultural field setting where multiple biotic or abiotic stresses may affect plants.

In this study, we investigated the effect of sequential and simultaneous inoculation of *P. infestans* and *A. solani in vitro* and *in planta* in both laboratory and field settings. To observe infections of both pathogens *in planta* at the microscopic level, we transformed *A. solani* with the green fluorescent protein (GFP) reporter, and co-inoculated potato with the *P. infestans* 88069 Td- tomato strain (McLellan et al., 2013) that expresses a red fluorescent protein reporter to distinguish each pathogen using confocal laser scanning microscopy.

## Materials and Methods

### *Phytophthora infestans* maintenance and inoculum preparation

*Phytophthora infestans* strains 88069 (Pieterse et al., 1991) and pink6 (Cooke et al., 2012) were maintained on Rye Sucrose Agar (RSA). *P. infestans* 88069 Td-tomato (expressing red fluorescence) (McLellan et al., 2013) was maintained on RSA supplemented with 5µg/ml geneticin (G418). Sporangium inoculum was prepared by flooding 14-day old cultures with 10 ml sterile tap water and gently rubbing the surface with a plastic L-shaped spatula and filtering the liquid through a 40 µM cell strainer to remove potential hyphae. The concentration of sporangia was determined using a Fuchs Rosenthal haemocytometer. Zoospore inoculum was prepared by flooding 14-day old cultures with 10 ml sterile 4°C tap water and to induce zoospore release the plates were subsequently placed at 4°C for 2-4 hours. The liquid was strained through 40 µm cell strainers and the concentration of zoospores was determined using a Fuchs Rosenthal haemocytometer.

### *Alternaria solani* maintenance and inoculum preparation

*Alternaria solani* strains AS112 (Odilbekov et al., 2014) and NL03003 (CBS 143772) (Iftikhar et al., 2017) were maintained on Potato Dextrose Agar (PDA) and V8 juice solid medium respectively. The plates were kept in the dark at RT (22°C) for 4 days and subsequently placed in an 18°C incubator equipped with UV-c light bulbs (model OSRAM HNS15G13 with dominant wavelength 254 nm) supplying 8 hours of UV-c light per day for 10 days to induce sporulation. *A*. s*olani* pCT74 GFP #2 was maintained on 20% PDA supplemented with 34µg/ml Hygromycin-B and kept in the dark at RT for 7 days. Multiple small agar plugs were subsequently transferred to sporulation medium (30 g CaCO_3_, 20 g Bacto agar, 1L MilliQ H_2_O, pH 7.4) (Shanin and Shepard, 1979) and covered with 2 ml sterile MilliQ water. The plates were incubated in the dark for 5 days at 18°C. Conidial inoculum was prepared by flooding the plates with 10 ml sterile tap water and gently rubbing of the growth using plastic L-shaped spatulas. The concentration of conidia was determined using a Fuchs Rosenthal haemocytometer.

### Transformation of *Alternaria solani*

Small agar plugs containing 7 days old *A. solani* NL03003 mycelium were cut from a PDA plate and inoculated in 15 ml PDB medium. The liquid culture was grown for 24 h on a rotary shaker (170 rpm) at 28°C and mycelium was collected from 2 ml of culture through centrifugation at 6000 x *g* for 10 min. The mycelium was washed twice with 0.7 M NaCl and collected again through centrifugation at 6000 x *g* for 10 min. 1 ml of kitalase solution (10 mg/ml in 0.7 M NaCl) was added to digest the mycelium for 2 h at 28°C, while shaking at 170 rpm. The resulting protoplasts were filtered through two layers of Miracloth and collected through centrifugation for 10 min at 700 x *g* at a temperature of 4°C. The protoplasts were washed in cold STC buffer (1 M Sorbitol, 50 mM Tris-HCL, pH 8.0, and 50 mM CaCl2) and finally aliquoted in STC buffer to obtain 5 million protoplasts in 70 μl buffer. 5 μg of pCT74 plasmid (10 μl) (Nova Lifetech Ltd, HK) (Lorang et al., 2001) was added to the protoplasts, gently mixed and the mix was placed on ice for 10 min. The sample was incubated for 5 min at 42°C and then placed back on ice for 10 min. 1.5 ml of 40% PEG 4.000 (in STC buffer and at room temperature, 22°C) was added and 400 μl of the transformed protoplasts were added to 50 ml molten (40-45°C) regeneration medium (5 g yeast extract, 5 g casamino acid, 1M sucrose and 0.8% agar). The regeneration medium was then divided over two deep petri dishes and incubated at room temperature (22°C) overnight. The next day, the plates were overlaid with 25 ml of PDA containing 34 μg/ml Hygromycin B. Emerging colonies were transferred to fresh PDA plates containing 34 μg/ml Hygromycin B. In total >15 transformants were obtained of which the most fluorescent ones were maintained.

### Plant material

*Solanum tuberosum* cv. Désirée was maintained in vitro in MS20 medium (Abreha et al., 2015). Two-week old *in vitro* plantlets were planted in 2L commercial soil (Exclusiv Blom & Plantjord, Emmaljunga Torvmull AB, Sweden) supplemented with 15 ml of “Osmocote exact 3-4 months” fertilizer beads (containing: Nitrogen- Phosphorus- Potassium 16-9-12+ 2 Magnesium oxide + trace elements). The plants were maintained in an artificial light plant chamber at 20°C and 65% RH, receiving 14 hours of 160 µmol/m^2^ light.

### Co- inoculation on solid medium

Two 5mm x 5mm agar plugs of 14 day old sporulating cultures of *P. infestans* 88069 and/ or *A. solani* NL03003 were inoculated onto V8 juice solid medium 3 cm apart. The prepared combinations were as follows: two *P. infestans* agar plugs, two *A. solani* agar plugs, or one *P. infestans* and one *A. solani* plug plated together. Additionally, plates containing only one agar plug were prepared for *A. solani* and *P. infestans*. Two sets of five plates per combination were prepared. One set was incubated at 20°C and one set at 18°C for 9 days (Figure 1A). The average radial growth diameter was determined every 3 days. The experiment was repeated 3 times. Statistical analysis comparing the radial growth was performed using one-way ANOVA followed by Fisher LSD pairwise comparison with 95% confidence in Minitab® (Version 18) Statistical Software package (Minitab Inc., 2010). Two separate ANOVAs were performed, one for the *P. infestans* growth and one for the *A. solani* growth.

**Figure 1.**
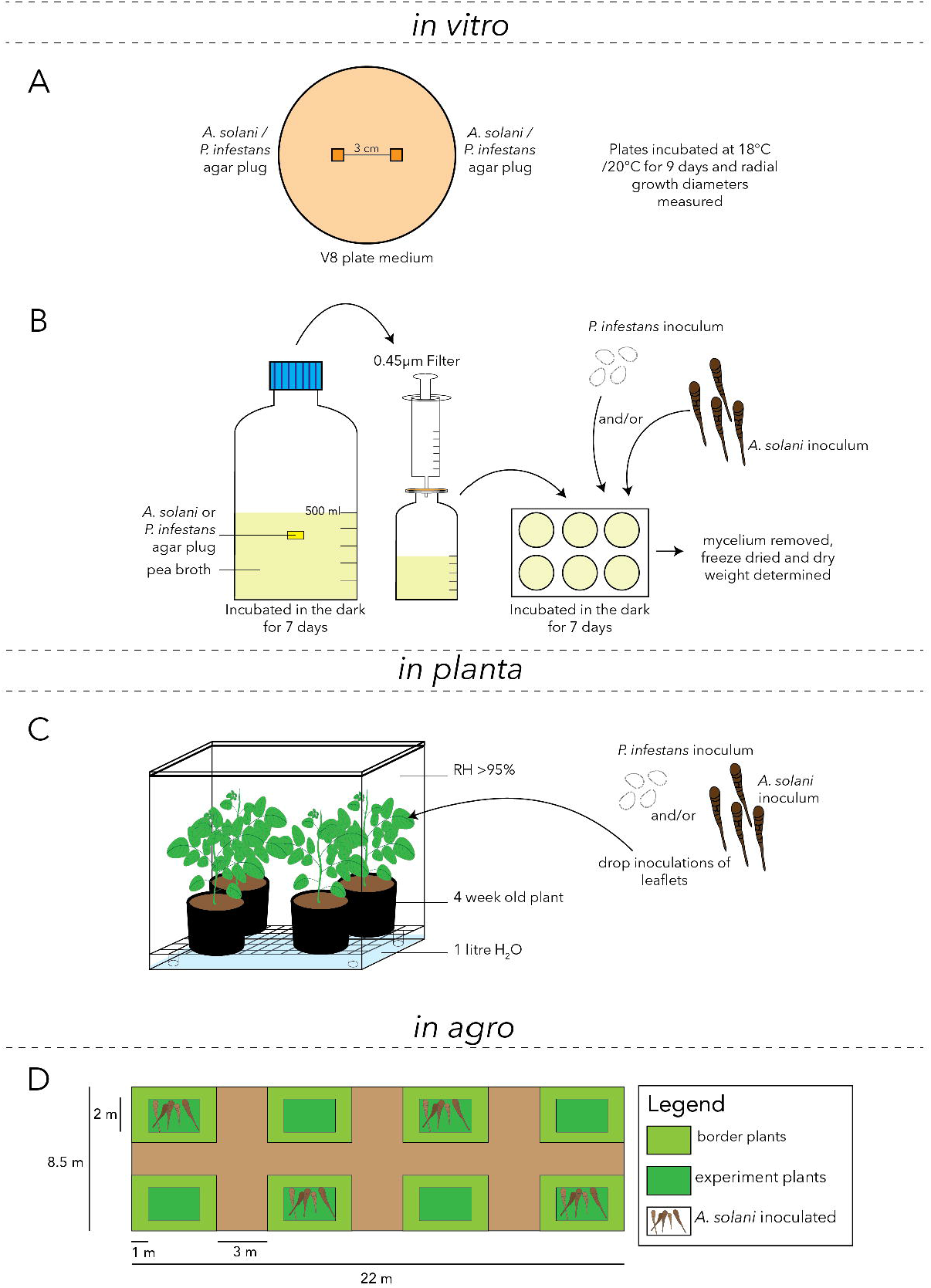
Schematic representation of the experimental set up and the field trial design used in this study. *In vitro* studies were performed on solid media (A) and in liquid cultures (B). Studies *in planta* were performed by inoculating whole plants contained in humidity boxes grown under controlled conditions in a growth chamber (C) and the field experiment *in agro* was performed according to the experimental plan depicted in (D).

### Co- inoculation in liquid medium

Two 10 mm x 15mm agar plugs of sporulating cultures of *P. infestans* 88069, *P. infestans* Pink6, *A. solani* NL03003, or *A. solani* NL03003 were added to 500 ml pea broth in 1L Duran bottles and incubated in the dark at RT (22°C) for 7 days. The *A. solani* liquid cultures were incubated on a shaking incubator at 50 rpm, whereas *P. infestans* cultures were incubated without shaking. After one week of growth the hyphae were removed and the broth was filter sterilized using 0.45 µm filters. 4 ml filtered medium was added to 6-well plates to which *P. infestans* inoculum (1 ml containing 10000 sporangia) was subsequently added. A further 5 ml of the filtered medium was added to 6-well plates to which *A. solani* inoculum (10 µl containing 100 conidia) was subsequently added. Filtered medium without previous pathogen growth was used as a control. Dual inoculation plates containing both pathogens were also prepared as described above. The cultures were incubated at RT in the dark for 7 days (Figure 1B). After 7 days, the mycelium was removed from the broth using an inoculation loop spatula and placed in pre-weighed 1.5 ml micro centrifuge tubes. The tubes were placed at −80°C overnight and subsequently lyophilized for 4 days. The dry weight was determined and statistical analysis comparing the dry weight was performed using one-way ANOVA followed by post hoc Tukey test. Separate ANOVAs were performed, one for the *P. infestans* dry weight and one for the *A. solani* dry weight. Statistical analysis comparing the *A. solani* dry weight and *P. infestans* plus *A. solani* dry weight, or the *P. infestans* dry weight and the *P. infestans* plus *A. solani* dry weight, were determined using two-sample t-tests. All statistical analyses were performed in Minitab® (Version 18) Statistical Software package (Minitab Inc., 2010). The experiment was repeated 3 times.

### Whole plant infections and lesion scoring

4-week old potato plants were placed in custom made acrylic boxes inside versatile environment incubators as described in Brouwer et al., 2020 (Figure 1C). From three fully expanded leaves in the middle of the plant canopy, top leaflets were drop-inoculated with two 10 µl droplets per plant, on either side of the mid vein. 25000 spores/ml *A. solani* NL03003 conidial suspension, 50000 *P. infestans* 88069 zoospores/ml or inoculum containing both 25000 *A. solani* conidia and 50000 *P. infestans* zoospores per ml was used. For the sequentially double inoculated plants, inoculation droplets containing one of the pathogens were placed on the leaflets and 24 hours later inoculation droplets containing the second pathogen were placed on the same spot. Per treatment, 24 leaflets were inoculated with two droplets, resulting in 48 lesion measurements per treatment. The lesion diameter for all plants was determined 5 days after the first inoculation. Statistical analysis comparing the lesion diameters was performed using one-way ANOVA followed by post hoc Tukey test in Minitab® (Version 18) Statistical Software package (Minitab Inc., 2010). The experiment was repeated 3 times.

### Confocal Laser Scanning Microscopy of pathogen infection

Small leaf punches containing the inoculation spot were cut and mounted on microscopy slides. When the inoculation droplet was still present no additional water was added, however, at the later time points when the inoculation droplet dried out sterile tap water was added on top of the leaf disc. All samples were analysed using a Zeiss LSM 880 (Zeiss, Germany) and excited with a 488 argon laser for GFP and a 561 nm DPSS laser for td-tomato RFP, collecting emissions between 493 and 556 nm for GFP, 566 and 637 nm for td-tomato RFP, as well as transmission data. All images were collected as 1024 x 1024 pixel frames. All images were processed in ZEN 3.1(blue edition) software (Zeiss, Germany) and ImageJ v1.50i (Schneider et al., 2012). The number of fluorescent pixels for GFP and RFP was determined in ImageJ v1.50i (Schneider et al., 2012) by loading the raw data CLSM files, splitting the channels, removing salt & pepper type background noise, binarizing the image and using the histogram list function to obtain the number of pixels containing fluorescence.

### Inoculated field experiment

A field trial was carried out in 2020 at Helgegården (coordinates 56.018043, 14.051363 DDM), about 5 km west of Kristianstad, Sweden. The trial was planted and managed by the Swedish Rural Economy and Agricultural Societies (Hush ållningssällskapet). The size of the trial was 8.5 by 22 m and it contained 8 smaller plots separated by 3 m soil borders (Figure 1D). The field was fertilized with nitrogen (N), phosphorous (P) and potassium (K) according to common practice for potato: 200, 80 and 215 kg/ha for N, P and K, respectively. *Solanum tuberosum* cv. Kuras was planted in mid-June. Each plot contained 5 rows of which the three in the middle were used in the experiment and the rows of plants on the sides were considered to be the border plants. These were not used for scoring disease symptoms. The trial had four replicate blocks. In each block one of the two plots was inoculated with *A. solani* AS112 infected barley kernels (Euroblight protocol: https://agro.au.dk/fileadmin/Alternaria_solani_field_inoculation_kernels_2015.pdf) (Figure 1D). Two applications of 30 grams of infected kernels were spread in each plot. The first inoculation occurred on July 15 and the second on August 5. All plants were treated weekly with late blight fungicides (Revus®, a.i. mandipropamid and Ranman Top®, a.i. cyazofamid, were alternated), with no or minor effect on early blight, until the emergence of clear early blight symptoms in the *A. solani* inoculated plots. The last late blight fungicide treatment was carried out on Aug 10. Eleven days after the application of the last late blight fungicide, all plots were inoculated with *P. infestans* from natural occurring infection from our other field trials in Mosslunda, (coordinates 55.98384, 14.10611 DDM) 5 km south of Kristianstad, Sweden. *P. infestans* inoculation was achieved by brushing infected material over the plants in each plot and leaving the infected material in between the border plants. The disease symptoms for both early blight and late blight were repeatedly scored for 5 weeks by visual inspection of lesion morphologies (see Figure 8A). The disease severity, expressed as the percentage of infection per plot, was determined using the methods described in Duarte et al., (2013) and Liljeroth et al., (2016) for early and late blight, respectively. The relative area under the disease progress curve (rAUDPC) during the assessment period, as described by Odilbekov (2020) was calculated for both late blight and early blight. The relation between late and early blight disease severity was investigated using regression analysis with Pearson correlation coefficient in Minitab® (Version 18) Statistical Software package (Minitab Inc., 2010).

## Results

### Field observations of early and late blight in the same field

In our potato field trials, natural infection by *P. infestans* infection is normally observed earlier in the season than that of *A. solani* (data not shown). When *P. infestans* is already present in a field, this does not appear to hinder subsequent *A. solani* infection. However, in 2017 *A. solani* infection occurred early and subsequently, *P. infestans* infection appeared, with late blight severity up to 0.1% observed. Despite the presence of *P. infestans* inoculum in the field, a few days later no late blight was visible anymore, yet the early blight severity reached above 20% (sFigure 1). Based on this field observation we hypothesized that *P. infestans* was prevented from infection either by a direct interaction with *A. solani*, by *A. solani* derived compounds, or by the plant response triggered by *A. solani* infection.

### Reduced *P. infestans* radial growth when co-plated with *A. solani*

In order to study whether *A. solani* and *P. infestans* growth on agar is affected by the presence of the other pathogen, a growth experiment of both pathogens cultured on the same media plate either alone, with a second plug of the same pathogen, or with the other pathogen was performed. The radial growth of *A. solani* was not influenced by the presence of *A. solani* or *P. infestans* (Figure 2). However, the radial growth of *P. infestans* was significantly negatively affected by the presence of *A. solani* on the same plate. This result was obtained both when the plates were incubated at 20°C (Figure 2) and 18°C (sFigure 2).

**Figure 2.**
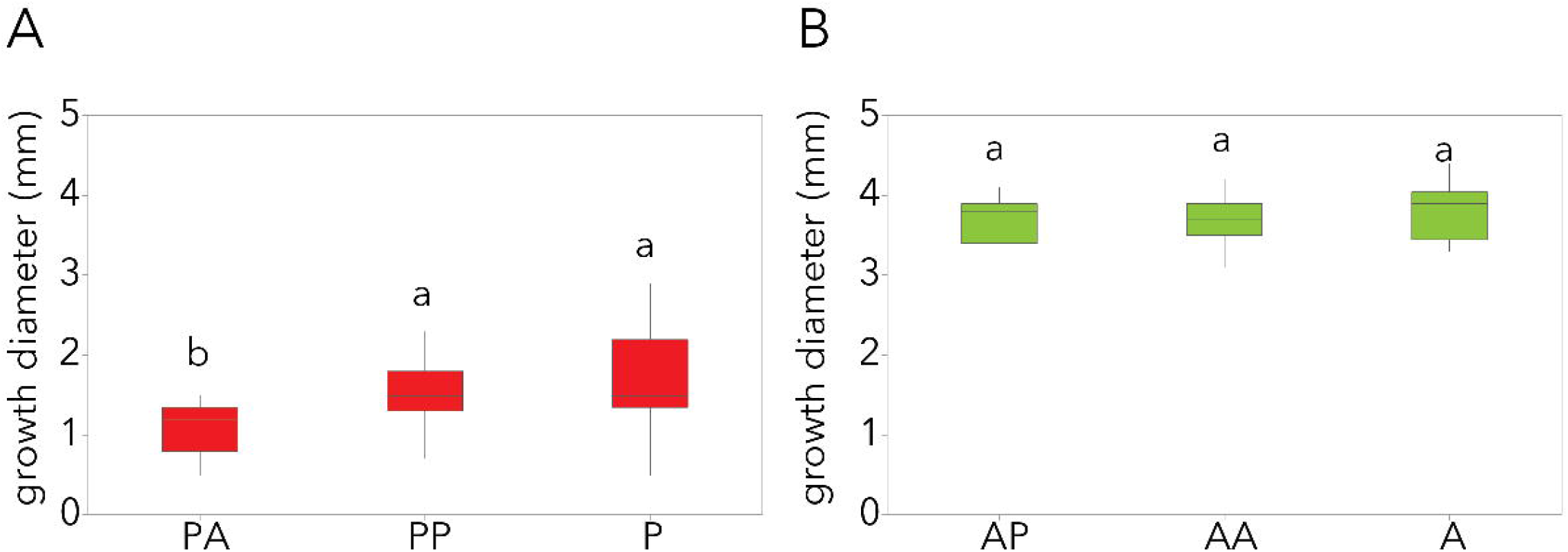
Radial growth after 9 days on V8 solid medium at 20°C of A) *P. infestans* 88069 (P) and B) *A. solani* NL03003 (A) either plated with each other (PA and AP), plated with themselves (PP and AA) or alone (P and A). N=13 and the letter labels correspond to the Fisher LSD grouping determined by one-way ANOVA followed by Fisher LSD pairwise comparison with 95% confidence. Two separate ANOVAs were performed, one for the *P. infestans* growth and one for the *A. solani* growth.

### Decreased growth of both pathogens in liquid medium that previously harboured *A. solani* growth

Since the growth of *P. infestans* was negatively affected by *A. solani* on solid medium, the influence of both pathogens on each other’s growth was further analysed in a liquid medium experiment. Both pathogens were grown in filtered control medium that previously contained *A. solani* or *P. infestans* cultures, respectively. Additionally, both pathogens were also added together to control liquid medium (Figure 3). When *P. infestans* was grown in medium that previously contained *A. solani* the dry weight of hyphae was significantly lower than when *P. infestans* was grown in control medium, or media that previously contained *P. infestans* either of the same strain or another strain (Figure 3A and D). The other way around, *A. solani* growth was also reduced compared to control medium when grown in medium that previously contained *P. infestans* or *A. solani* of the same strain and even more when grown in medium that previously harboured another strain of *A. solani* (Figure 3B). However, these reductions were much smaller than the reduction observed for *P*. i*nfestans* grown in medium that previously harboured *A. solani*. When the pathogens were added together in control medium, growth visually looked like *A. solani* growth (sFigure 3). However, the resulting dry weight was significantly higher than the growth of *A. solani* alone in control medium (p = 0.026) (Figure 3C).

**Figure 3.**
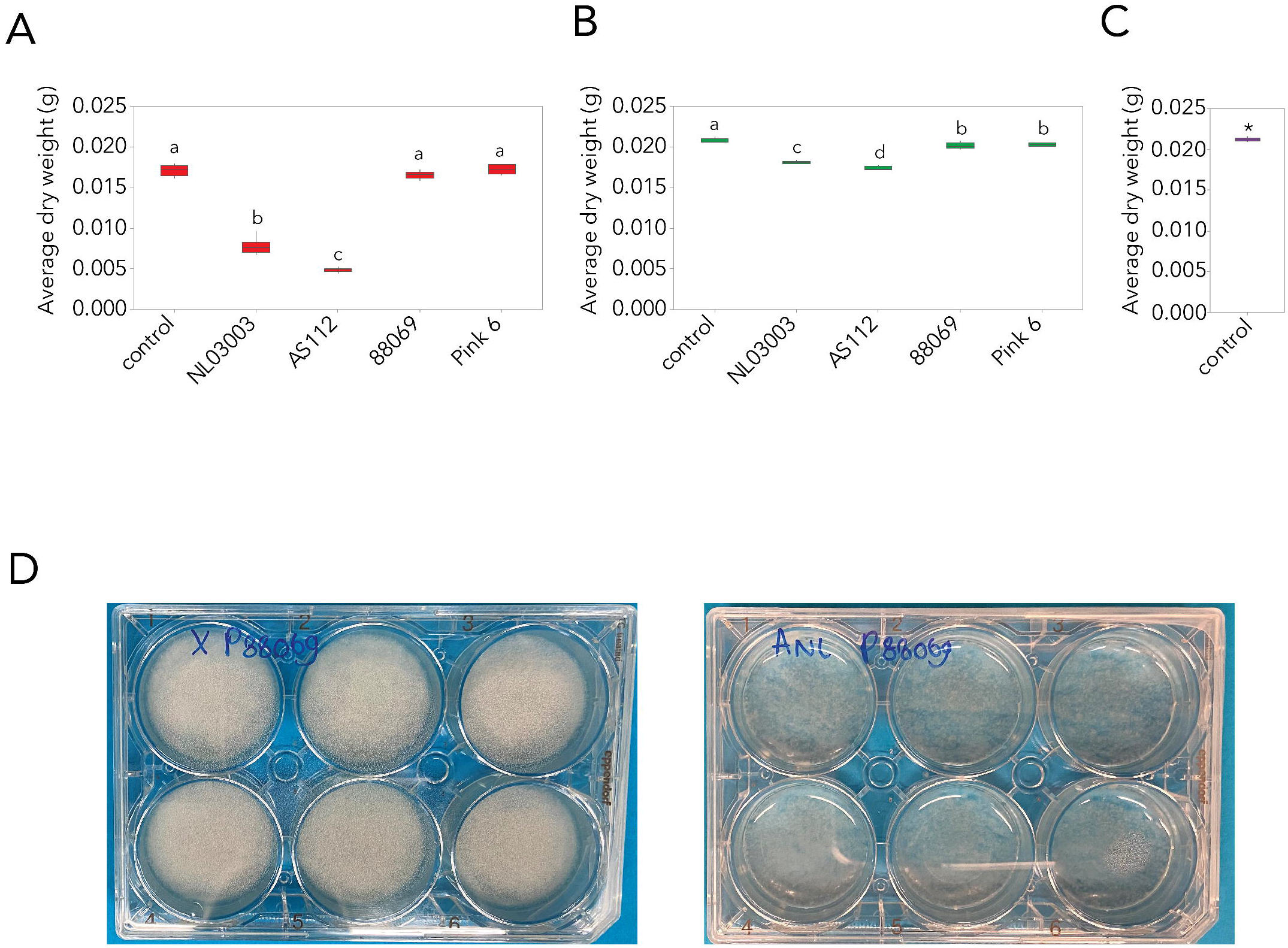
Average dry weight of produced hyphae in different liquid medium after 7 days. A) Dry weight of hyphae of *P. infestans* 88069 grown in control medium or medium that previously contained *A. solani* NL03003, *A. solani* AS112, *P. infestans* 88069, or *P. infestans* Pink 6. B) Dry weight of hyphae of A. solani AS112 grown in control or medium that previously contained *A. solani* NL03003, *A. solani* AS112, *P. infestans* 88069, or *P. infestans Pink 6*. C) Dry weight of hyphae of *P. infestans* 88069 and *A. solani* AS112 co-inoculated in control medium. All media was filtered through 0.45µM filters before addition of either *P. infestans* 88069 or A. solani AS112 spores. N=6 and the letters represent the grouping based on One-Way ANOVA followed by Tukey post hoc test. Two separate ANOVAs were performed, one for the *P. infestans* dry weight and one for the *A. solani* dry weight. Asterisk indicates the significant difference between growth in graph C and the growth of *A. solani* in control medium in graph B. D) Photographs of the growth of *P. infestans* 88069 in control medium (left) and medium previously harbouring *A. solani* NL03003 (right).

### *P. infestans* sporangial and hyphal tip leakage in liquid that previously harboured *A. solani*

In order to determine whether the decrease in growth of *P. infestans* in liquid medium that previously harboured *A. solani* was caused by delayed germination or something else, the growth progression was visualised using inverted bright field microscopy (Figure 4). During the first 6 hours of inoculation the sporangia started to germinate, however, a significant proportion of the sporangia displayed bursting on the side and subsequent leakage of sporangial cytoplasm (Figure 4A). Additionally, for the sporangia that germinated, swelling and subsequent leakage of cytoplasm from the hyphal tip was also observed (Figure 4B). Eight days after inoculation, some *P. infestans* growth occurred, but many of the sporangia displayed the burst phenotype and never germinated. Additionally, the hyphal growth that was established sometimes displayed hyphal tip bursting. The bursting of sporangia and hyphal tips was not observed for *P. infestans* grown in control medium or medium that previously harboured *P. infestans* growth.

**Figure 4.**
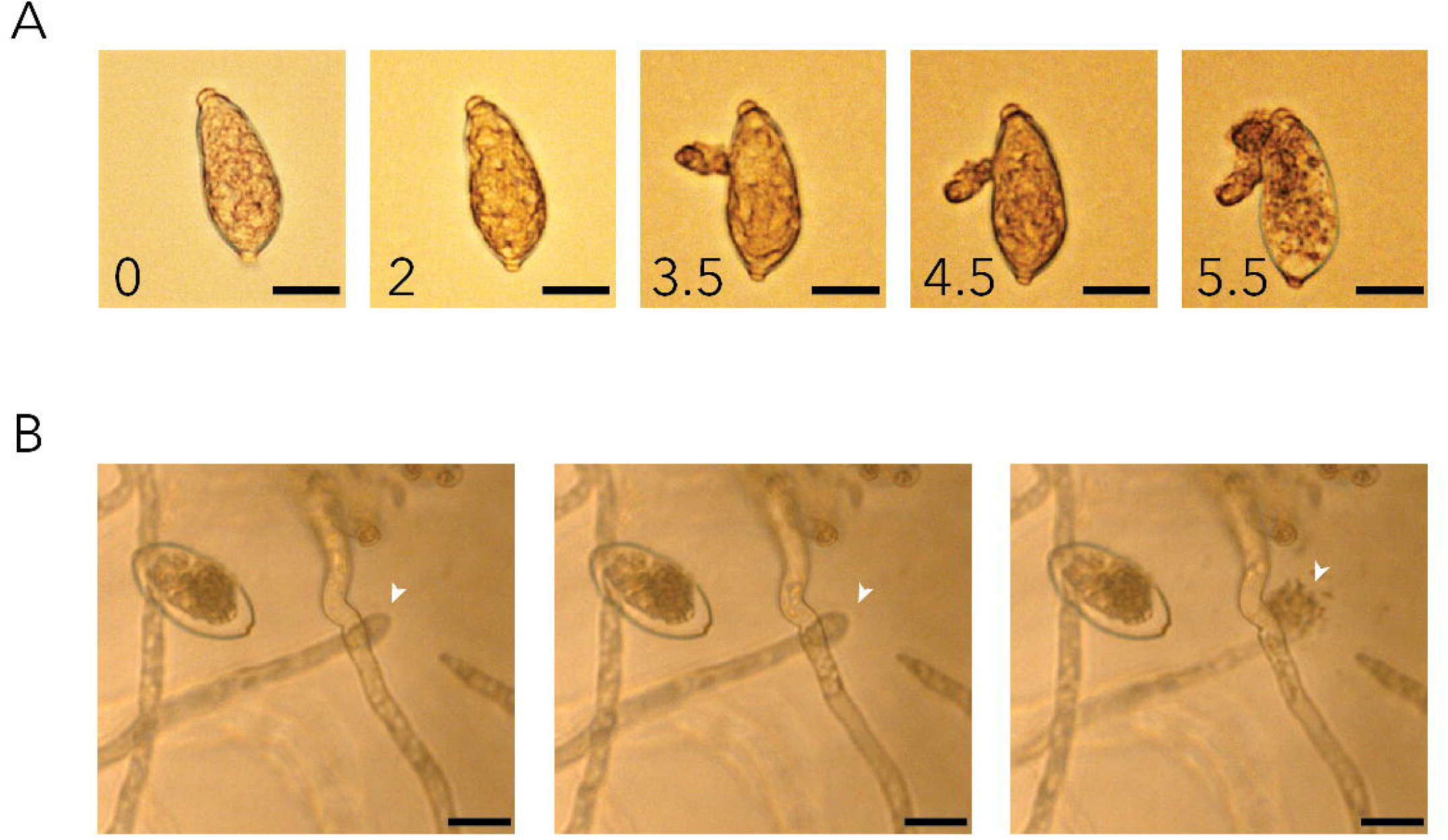
Micrographs of growth of *P. infestans* 88069 in liquid medium that previously contained *A. solani* NL03003. The black scale bar for all images corresponds to 20µm A) Sporangium at 0, 2, 3.5, 4.5, and 5.5 hours post inoculation in medium. B) Bursting of hyphal tips (indicated with arrow) observed 4 days post inoculation. The displayed images were collected with a time interval of 30 minutes.

### Simultaneous co-infection *in planta* results in larger necrotic lesions

With the aim of studying the effect of co-infection of potato leaflets with both pathogens, a growth chamber bioassay with simultaneous and sequential inoculation was performed. The lesion size was determined 5 days post inoculation for leaflets inoculated with either *P. infestans, A. solani*, both *P. infestans* and *A. solani* simultaneously, *P. infestans* followed by *A. solani* 24 hour later or *A. solani* followed by *P. infestans* 24 hours later (Figure 5). The simultaneous inoculation of both pathogens in the same spot resulted in the largest lesions that were necrotic and resembled *A. solani* lesions (Figure 5B). When adding the other pathogen 24 hours after inoculation with the first pathogen, the addition of *P. infestans* did not result in a difference in lesion size compared to only *A. solani* inoculation (Figure 5B). However, the addition of *A. solani* 24 hours after *P. infestans*, resulted in a significant increase in lesion size compared to *P. infestans* only inoculated leaflets (Figure 5B).

**Figure 5.**
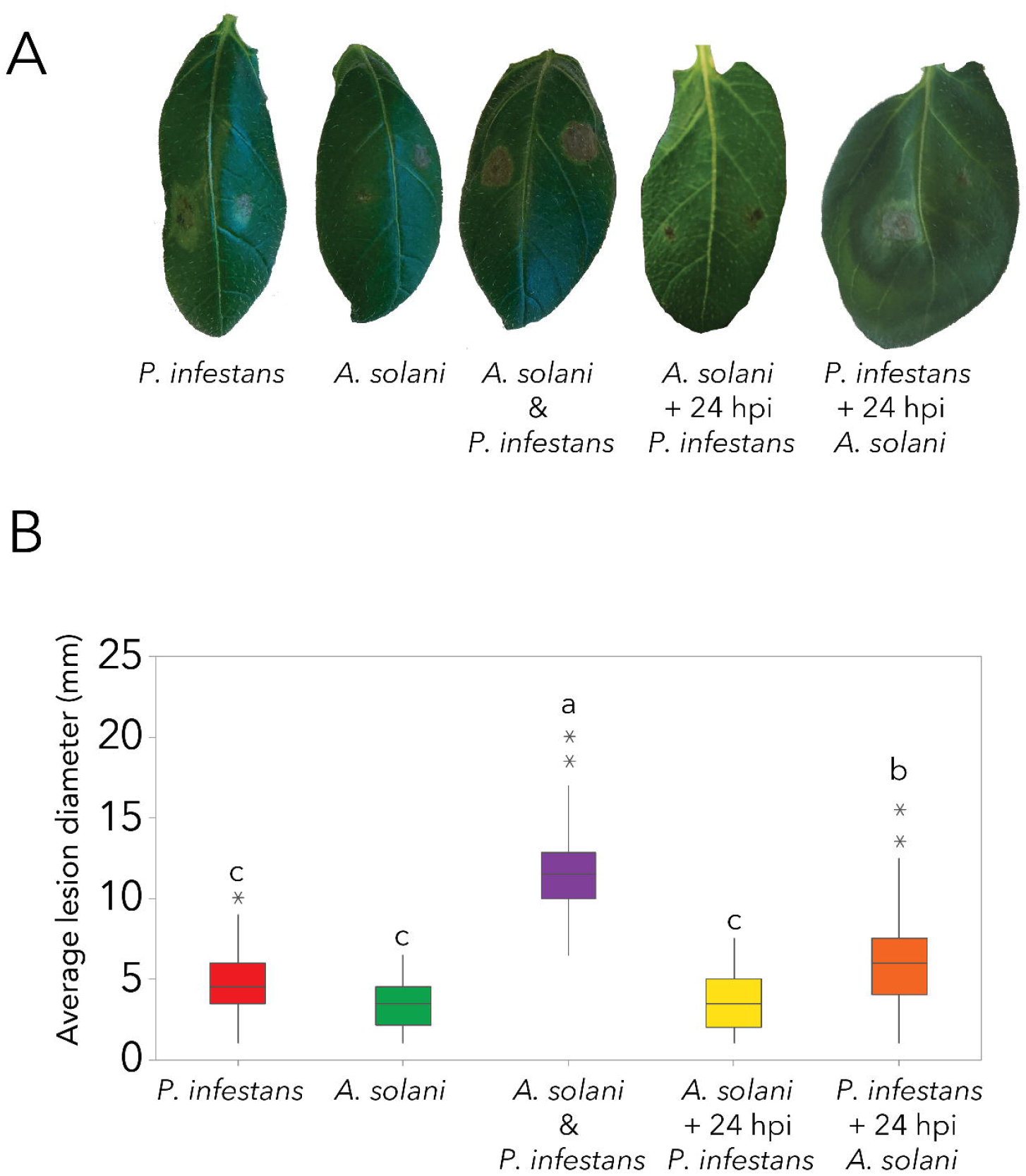
Co- and sequential inoculations of potato leaflets with *P. infestans* and *A. solani*. A) Representative photographs B) Average lesion diameter at 5 dpi for the plant inoculated with *P. infestans, A. solani, P. infestans* + *A. solani, A*. solani followed by *P. infestans* 24 hours later and *A. solani* followed by *P. infestans* 24 hours later. N=48 and the letters represent the grouping based on One-Way ANOVA followed by Tukey post hoc test.

### Microscopy of simultaneous and sequential inoculation with *A. solani* and *P. infestans*

Using the same inoculation conditions for which the lesion size was determined, the infection process of drop inoculations on whole plants on the adaxial side of leaves was followed and imaged. Using confocal laser scanning microscopy (CLSM) colonization of the mesophyll and emergence through the stomata at the abaxial surface was determined. In the single inoculated conditions, germ tube formation was observed at 1 hpi and colonization between the mesophyll cells by *P. infestans* was observed at 24hpi and continued further (Figure 6AB). Germinating *A. solani* conidia resulted in extensive growth in both directions of the conidia, both towards and away from the leaf surface (Figure 6C). The extensive growth of *A. solani* on top of the epidermal cells in the simultaneous co-inoculated conditions, to a large extend obstructed scanning of tissue layers underneath the network of hyphae. However, imaging at 48hpi of the abaxial side of the leaves revealed colonization of the mesophyll cell layers by *A. solani* (Figure 6D).

**Figure 6.**
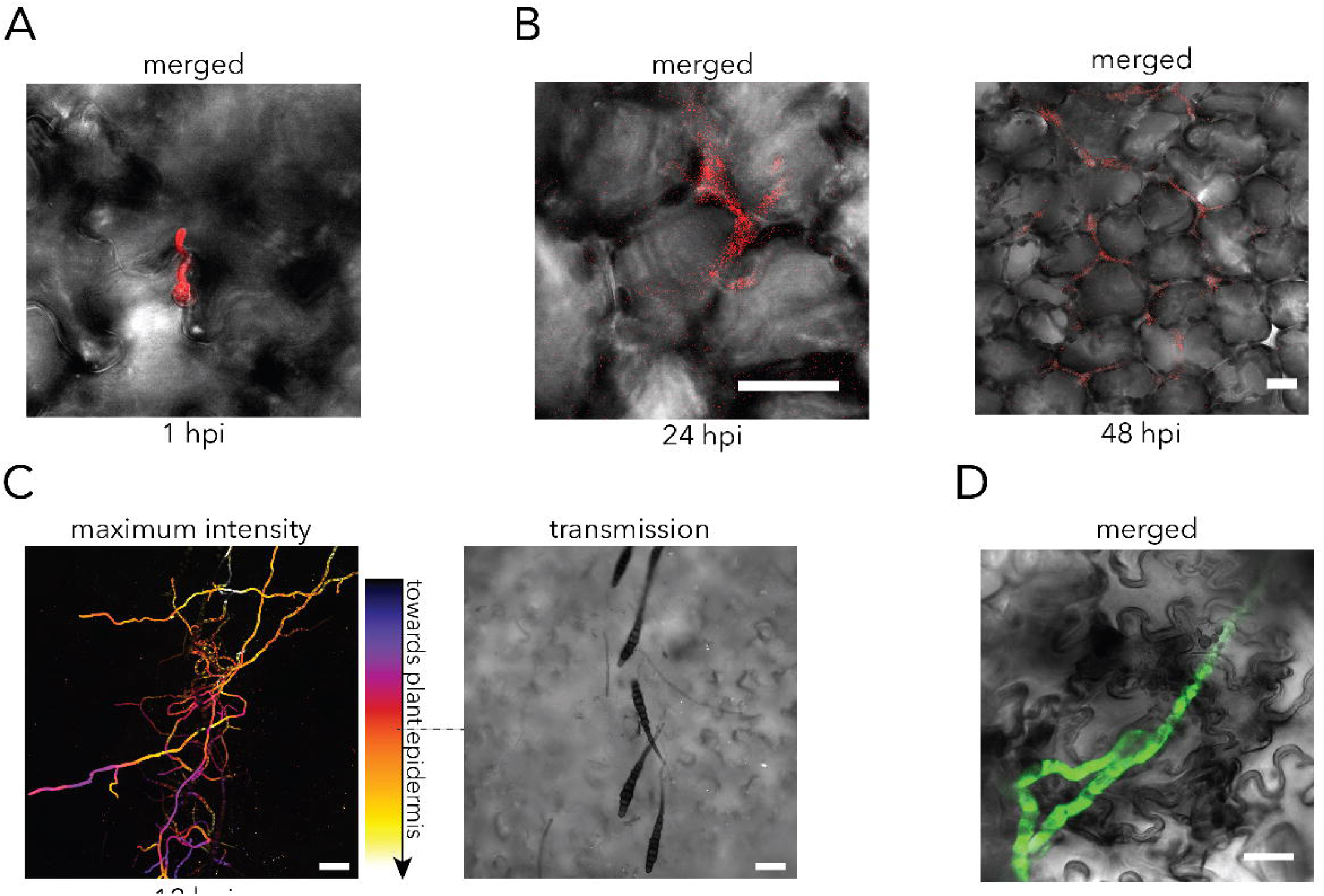
Representative Confocal Laser Scanning Micrographs obtained of single infection of potato leaves inoculated with *A. solani* or *P. infestans* on the adaxial side of the leaf. A) Germinated *P. infestans* zoospore. Scale bar corresponds to 50 µm B) *P. infestans* colonization within the mesophyll cell layer observed at 24 hpi and 48 hpi. Merged FRP and transmission micrographs of *P. infestans* td-tomato. Scale bars correspond to 20 µm C) Depth colour coded maximum intensity projection micrograph of germinating *A. solani* conidia, displaying growth both in the direction towards (white) the plant cells and away (purple) from the plant cells at 12 hpi in a leaf inoculated with *A. solani*. Transmission micrograph displayed from the middle of the z-stack shows the orientation of the *A. solani* conidia. Scale bar corresponds to 50 µm D) Merged GFP and transmission micrographs of *A. solani* GFP growth in green and the transmission channel, displaying emergence of *A. solani* from the plant mesophyll on the abaxial side of an *A. solani* inoculated leaf at 48 hpi. Scale bar corresponds to 20 µm

In the simultaneously co-inoculated samples, even though germination of *P*. infestans cysts and/or sporangia was initially observed, colonization of the potato leaf was rarely observed (Figure 7A). When sequential inoculation of *A. solani* and *P. infestans* was performed with 24 hours in between the inoculations, established *P. infestans* infection did not inhibit *A. solani* growth. After 48 hours, colonization of both *P. infestans* and *A. solani* was observed on the abaxial side of the leaves (Figure 7B). Contrastingly, when *A. solani* inoculation was followed with with *P. infestans* inoculation 24 hours later, only *A. solani* colonization of the leaf was observed (Figure 7C). However, auto fluorescence of necrotic cells, especially epidermal cells, was visible in the RFP channel (Figure 7C and sFigure 4).

**Figure 7.**
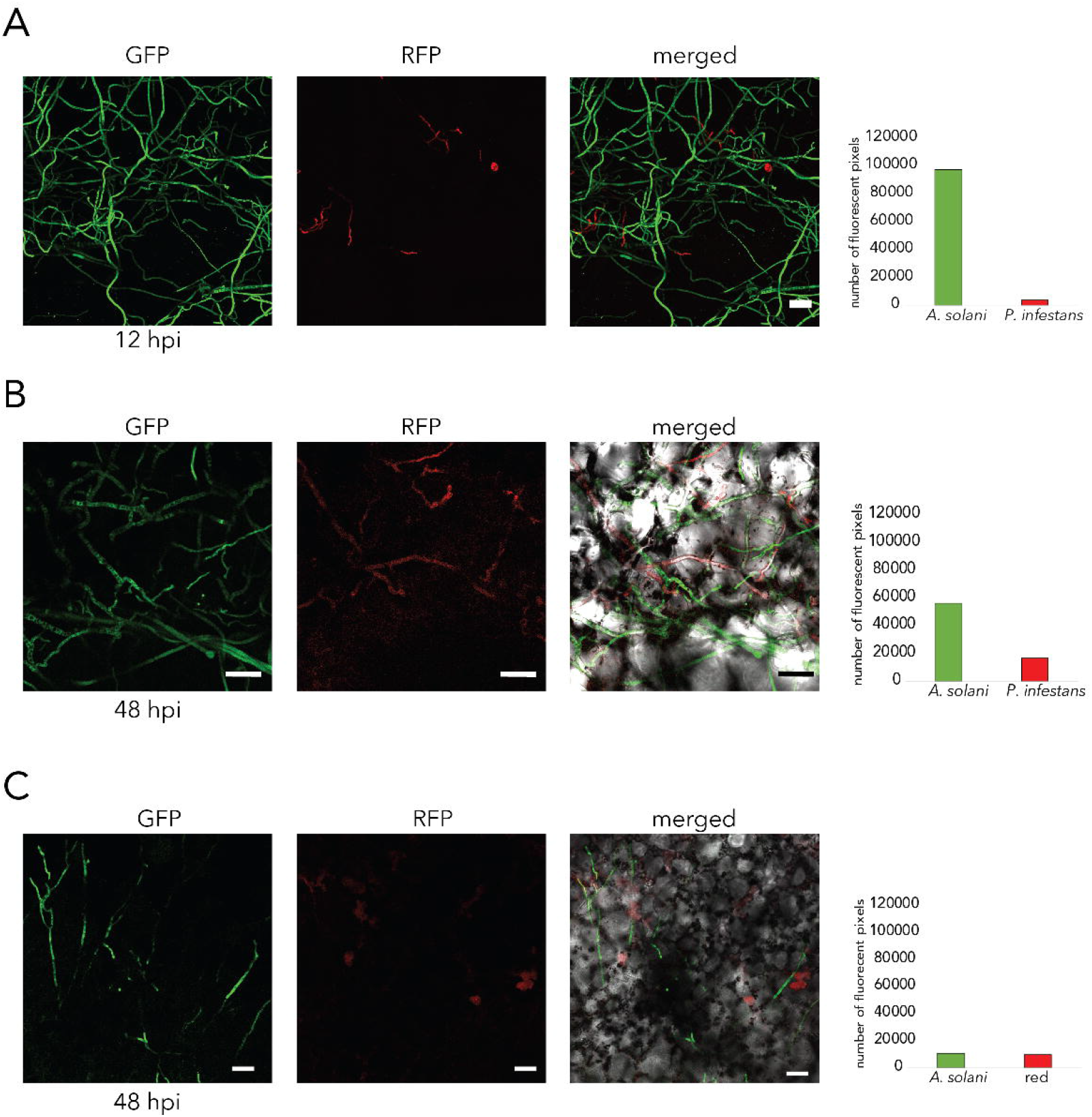
Representative Confocal Laser Scanning Micrographs obtained of infection of potato leaves inoculated with *A. solani* and *P. infestans* on the adaxial side of the leaf. Micrographs for *A. solani* GFP (GFP), *P. infestans* td-tomato (RFP), and merged image including the transmission channel for B and C. All scale bars correspond to 50 µm. Bar graphs indicate number of pixels containing GFP (*A. solani*) and RFP (*P. infestans* or C) autofluorescence from death cells) signals. A) Maximum intensity projection micrograph of simultaneous inoculation of *A. solani* and *P. infestans* imaged at 12 hpi B) Micrographs of co-inoculation with *P. infestans* followed by *A. solani* 24 hours later, imaged on the abaxial side 48 hpi with *A. solani*. F) Micrographs of co-inoculation with *A. solani* followed by *P. infestans* 24 hours later, imaged on the abaxial side of the leaf 48 hpi with *P. infestans*.

### The presence of *A. solani* limits establishment of *P. infestans* under field conditions

To determine whether the observed results from the growth chamber bio assays also apply *in agro*, (i.e. under agriculturally relevant conditions), an inoculated field experiment was performed (Figure 8). Potato cv. Kuras, was planted in mid-June and plants in half of the test plots were inoculated with *A. solani* AS112, in mid-July and early August, whilst also being treated with fungicides to prevent natural *P. infestans* infection. When clear early blight symptoms appeared in the *A. solani* inoculated blocks, all the blocks were inoculated with *P. infestans*. The first early blight symptoms appeared in the *A. solani* inoculated plots in the beginning of August and then the infection increased gradually with time. Later *A. solani* also spread in the direction of the prevailing wind, to the non-inoculated control plots. However, the rate of early blight infection was lower in all the non-inoculated plots. In one of the non-inoculated plots, away from the prevailing wind, the *A. solani* infection rate stayed much lower than in the other plots. On the 24^th^ of August the *A. solani* disease scores in inoculated plots varied between 5 and 8 % infection per plot, while in the non-inoculated control plots the *A. solani* disease scores ranged from 0.1 to 1.5%. After inoculation with *P. infestans* the disease symptoms of both diseases (Figure 8A) were scored over five weeks, to determine the influence of *A. solani* presence in the field on the occurrence of *P. infestans* induced late blight. In general, the late blight disease development rate was slower in plots inoculated with *A. solani* (Figure 8C). The presence of *A. solani* was shown to limit the development of late blight disease, over the whole assessment period, since the relative area under the disease progress curve (rAUDPC) of late blight was negatively correlated with the rAUDPC of early blight (R^2^=0.7865, p=0.003) (Figure 8D). Additionally, the percentage of late blight disease present at the end of the assessment period was highly significantly negatively correlated (R^2^=0.9199, p=0.00017) with the percentage of early blight present (Figure 8E).

**Figure 8.**
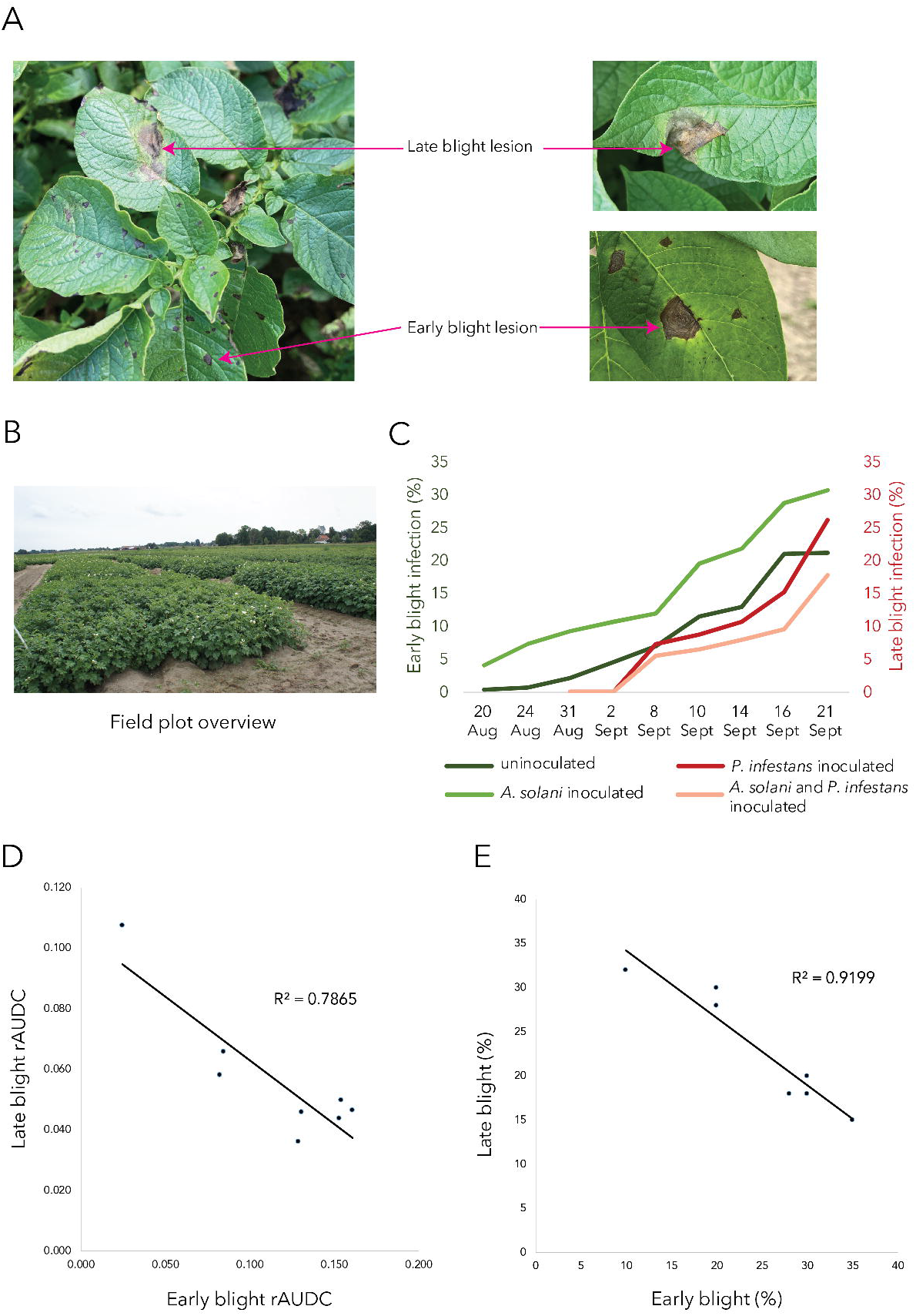
Field experiment on the effect of *A. solani* infection on development of *P. infestans*. A) Photographs of characteristic early and late blight lesions B) photograph from the field trial showing the field design C) Percentage of early and late blight during the experiment assessment period D) Relative area under the disease curve (rAUDC) for both late blight and early blight plotted against each other. Linear regression line displayed (R^2^=0.7865, p=0.003) Late blight and early blight disease percentage on the last scoring date (Sept 21) plotted against each other. Linear regression line displayed (R^2^=0.9199, p=0.00017).

## Discussion

In an agricultural setting, plants and plant pathogen interactions do not occur as simple binary interactions. An agricultural field contains a plethora of microbial species, both beneficial and pathogenic. However, many plant pathogenic species and their host plant interactions are studied as binary interactions. In this study, we investigated the interactions of two common pathogens of potato from *in vitro* interactions, to *in agro* interactions. We discovered that the late blight pathogen *P. infestans* is limited in growth in the presence of the early blight pathogen *A. solani* both *in vitro* and *in planta* depending on which pathogen is present first. *In agro*, the establishment of *A. solani* infections in a field limits the potential for *P. infestans* to establish subsequent infections.

The *in vitro* growth rate of *P. infestans* on agar plate medium was reduced when it was plated together with *A. solani*. Since this *A. solani* isolate has a faster overall growth rate, it is difficult to determine whether the reduction in growth of *P. infestans* was due to a space limitation on the plate medium or other competing factors, such as the production of antagonistic metabolites by *A. solani*. To understand this relationship further, we grew each pathogen in a liquid culture system, using media that previously contained the other pathogen. Thus, we were able to measure the growth of each pathogen individually. When growing *P. infestans* in medium that previously contained *A. solani* we observed cytoplasmic leakage from both sporangia and hyphae. *Alternaria* species are known to produce secondary metabolites toxic to both plants, animals, and humans (Meena and Samal, 2019, Gotthardt et al., 2019). Several host specific toxins have been characterized and described for plant disease causing *Alternaria* species, and though the site of action differs for these toxins, their aim is to trigger cell death in the host (Meena and Samaal, 2019). Yet, there is limited knowledge on the specific toxins produced by *A. solani* and the toxin targets. Additionally to host cell death, mycotoxins of phytopathogenic fungi are reportedly able to target other phytopathogenic species (Venkatesh and Keller, 2019), e.g. the maize pathogens *Ustilago maydis* and *Fusarium verticilliodes*. Jonkers et al. (2012), showed that *F. verticilliodes* secretes toxins and increases expression of cell wall degrading enzymes when grown together in liquid with *U. maydis*. After initial accumulation of biomass for both species when grown together, the biomass of *U. maydis* declined most severely over time, indicating efficacy of the *F. verticilliodes* toxins on *U. maydis*. It is possible that the reduction in *P. infestans* growth and the bursting phenotype we observed, is due to mycotoxins produced by *A. solani*.

Simultaneous co-inoculation of *P. infestans* and *A. solani in planta* resulted in larger necrotic lesions than single inoculation of either *P. infestans* or *A. solani*. Microscopic analysis revealed that simultaneously co-inoculated lesions mainly displayed successful colonization of the host tissue by *A. solani*. Although, *P. infestans* spores germinated, established infection of the host tissue was rarely observed. Yet the macroscopic lesions observed in the simultaneous co-inoculation were larger than the lesions of *A. solani* inoculation only. The presence of *P. infestans* thus seems to somehow enhance the virulence of, or promote host susceptibility to, *A. solani*. It is possible that *P. infestans* has a direct positive effect on *A. solani*, or, alternatively, that the presence of *P. infestans* renders the host more susceptible to infection by *A. solani*. In a previous study, we observed similar larger lesion development in salicylic acid (SA) deficient *NahG* plant lines (Brouwer et al., 2020). This indicates a role of intact SA signalling for potato defences against *A. solani*, during the biotrophic phase of *P. infestans* infection levels of SA increase (Zhou et al., 2018). Interestingly, exogenous application of SA to detached tomato leaves rendered the leaves more susceptible to *A. solani* with increased lesions compared SA untreated plants (Rahman et al., 2012). Contrastingly, exogenous application of SA to whole tomato plants grown in hydroponics resulted in increased *A. solani* resistance (Spletzer and Enyedi, 1999). These experiments indicate different roles of SA in defence against *A. solani*, increased susceptibility locally and induced resistance by systemically acquired resistance signals. The co-inoculation of both *A. solani* and *P. infestans* at the same spot on the leaf might result in local increased levels of SA that render the plants more susceptible to *A. solani* infection and result in the observed larges lesions. Additionally, *A. solani* could benefit from the plethora of *P. infestans* secreted effectors that manipulate host immune responses, along with cell wall degrading enzymes and other secreted enzymes such as isochorismatases that could interfere with SA signalling (Leesutthiphonchai et al., 2018). Even though *P. infestans* rarely established infection of the host tissue in the co- inoculated samples, it was previously shown that pre-infection germinated cysts and *in vitro* created appressoria already secrete proteins such as RXLR effectors and cell wall degrading enzymes (Resjö et al., 2017). Thus this indicates that even the presence of germinated *P. infestans* could provide effectors and enzymes that could potentially aid the infection of *A. solani*.

*A. solani* was able to colonise potato when it was inoculated *in planta* at the same site used to inoculate *P. infestans* 24 hours earlier. Larger lesions were observed when *A. solani* was inoculated after *P. infestans*, compared to the controls where *A. solani* was inoculated after *A. solani*. However, when *P infestans* was inoculated 24 hours after *A. solani*, this did not significantly change the lesion size. Microscopic analysis revealed that the germination of *P. infestans* in the presence of *A. solani* wasn’t inhibited, yet a network of fungal hyphae was observed in which the *P. infestans* germinated spores appeared entangled. It is thus possible, that the decrease in successful colonization by *P. infestans* is due to the formation of a physical barrier in the form of *A. solani* hyphae. However, the *in vitro* experiments indicate the potential for a direct effect of *A. solani* metabolites or secreted proteins such as antimicrobial effectors on *P. infestans*. Additionally, the establishment of *A. solani* infection in the host could trigger plant responses that negatively influence the potential for *P. infestans* to establish infection. *P. infestans* establishes successful infection of potato by employing effectors to suppress host immune responses but also to limit the damage to the plant, since it requires living tissue for its biotrophic life stage (Leesutthiphonchai et al., 2018). The necrotroph *A. solani*, however, triggers cell death to acquire nutrients. When *A. solani* is inoculated before *P. infestans* this might thus ensure the conditions required for the biotrophic phase of the *P. infestans* infection are not met, and hence the successful colonization by *P. infestans* is blocked.

In line with the results obtained for the sequential inoculation of whole plants in the laboratory setting, in the field we observed a reduction of late blight symptoms when the establishment of early blight occurred before the inoculation with *P. infestans*. In the laboratory experiments both pathogens were inoculated at the same spot, therefore a physical effect of the presence of the *A. solani* growth or triggered cell death could not be excluded. In the field experiment, however, the inoculation spots on the leaves were not controlled but occurred in a similar way to natural infections. Yet, the presence of early blight caused by *A. solani* had a negative effect on the establishment of late blight caused by *P. infestans*. This was found both in an observation of a late blight trial that unintentionally became infected with *A. solani* and later in a designed field trial where plots were inoculated first with *A. solani* and later with *P. infestans*. The results indicate that the negative effect of *A. solani* on the ability of *P. infestans* to infect cannot solely be explained by a physical barrier but likely also involves antagonistic metabolites and induction or hampered suppression of defence responses in the host plant.

In order to successfully continue protection of food production in the future, new knowledge to improve our understanding of plant pathogen interactions, especially in agriculturally relevant settings is required. In this study we showed that the disease-causing efficiency of the late blight pathogen is negatively correlated with the presence of early blight in the same field. Multipartite interactions between host and multiple pathogens such as those observed in this study present interesting opportunities to study the direct and indirect interactions of multiple organisms that result in differences in disease establishment and severity. These data can also be used towards the development of control strategies that exploit the plant responses triggered by one pathogen in combatting another pathogen. Additionally, compounds e.g. toxins and proteins secreted by one pathogen might be utilized as an ‘antibiotic’ against another pathogen or at least give clues to targets for novel control agents. To gain a deeper understanding of the interactions in this study, careful analyses of gene expression changes and metabolomics in multipartite interactions are required. Of special interest would be the study of multipartite interactions further under agriculturally relevant field conditions.

## Acknowledgements

We would like to thank Maja Brus-Skalej and Linnea Almqvist Stridh for technical assistance in the field experiment. This work was supported by grant 2019/00881 from the Swedish research council Formas (to LGB) and the field trial was supported by grant PA1169/18 (to EL) from Partership Alnarp. This project has also received funding from the European Union’s Horizon 2020 research and innovation programme under Grant Agreement No 774340 (Organic Plus).

## Data sharing statement

The data that support the findings of this study are available from the corresponding author upon reasonable request.

**sFigure 1** Field scoring of late blight severity and early blight severity in late blight trial in Hovby, Sweden in 2017. Early blight severity was initially not scored, but was started once no late blight was detected after the 22^nd^ of August. The disappearance of late blight after the 22 of August is likely due to the heavy early blight infection, limiting the visibility of very low levels of *P. infestans*.

**sFigure 2** Radial growth after 9 days on V8 medium at 18°C of both *P. infestans* 88069 (P) and *A. solani* NL03003 (A) either plated with each other (PA and AP), plated with themselves (PP and AA) or by itself (P and A). N=13 and the letter labels correspond to the Fisher LSD grouping determined by one-way ANOVA followed by Fisher LSD pairwise comparison with 95% confidence. Two separate ANOVAs were performed, one for the *P. infestans* growth and one for the *A. solani* growth.

**sFigure 3** Photograph of growth after 7 days of *P. infestans* 88069 and *A. solani* AS112 co-inoculation in control medium

**sFigure 4** Autofluorescence in the RFP channel of necrotic epidermal cells on the abaxial side in leaf in sample inoculated with *A. solani* and 24 hpi later with *P. infestans*. Same area of the leaf as imaged in Figure 7C

