## Supplementary figures and images for "Double trouble or a blessing in disguise? Co-infection of potato with the causal agents of late and early blight"

### Supplemental Figure 1

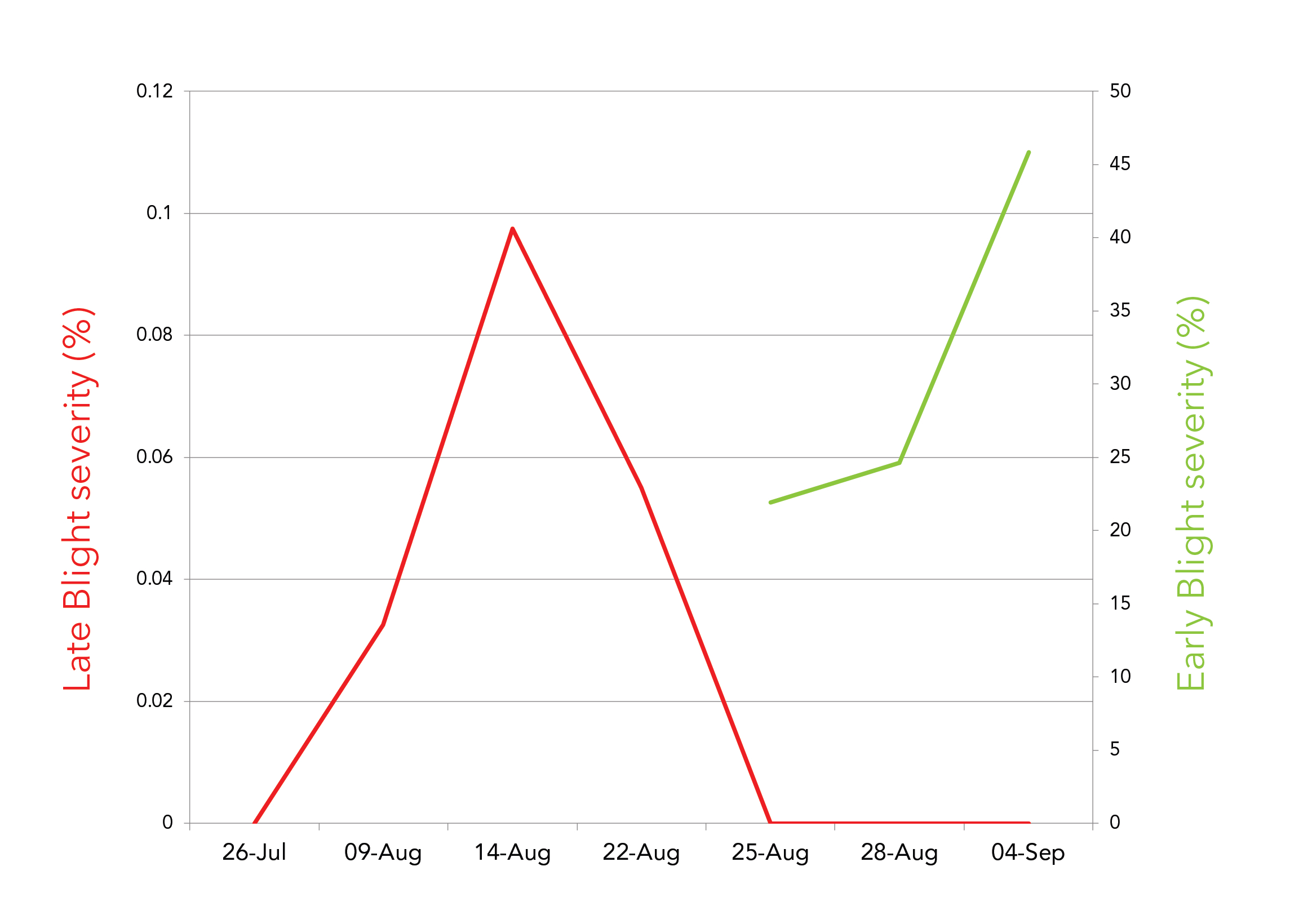

### Supplemental Figure 2

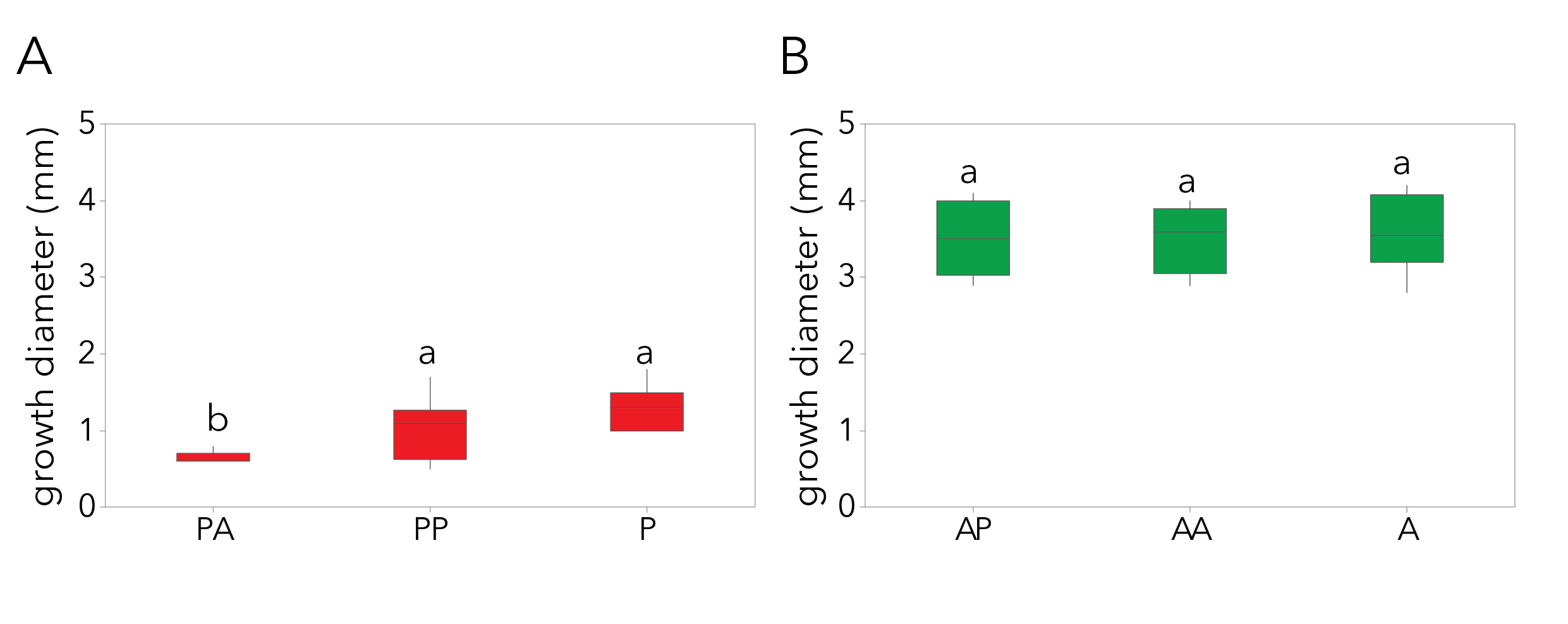

### Supplemental Figure 3

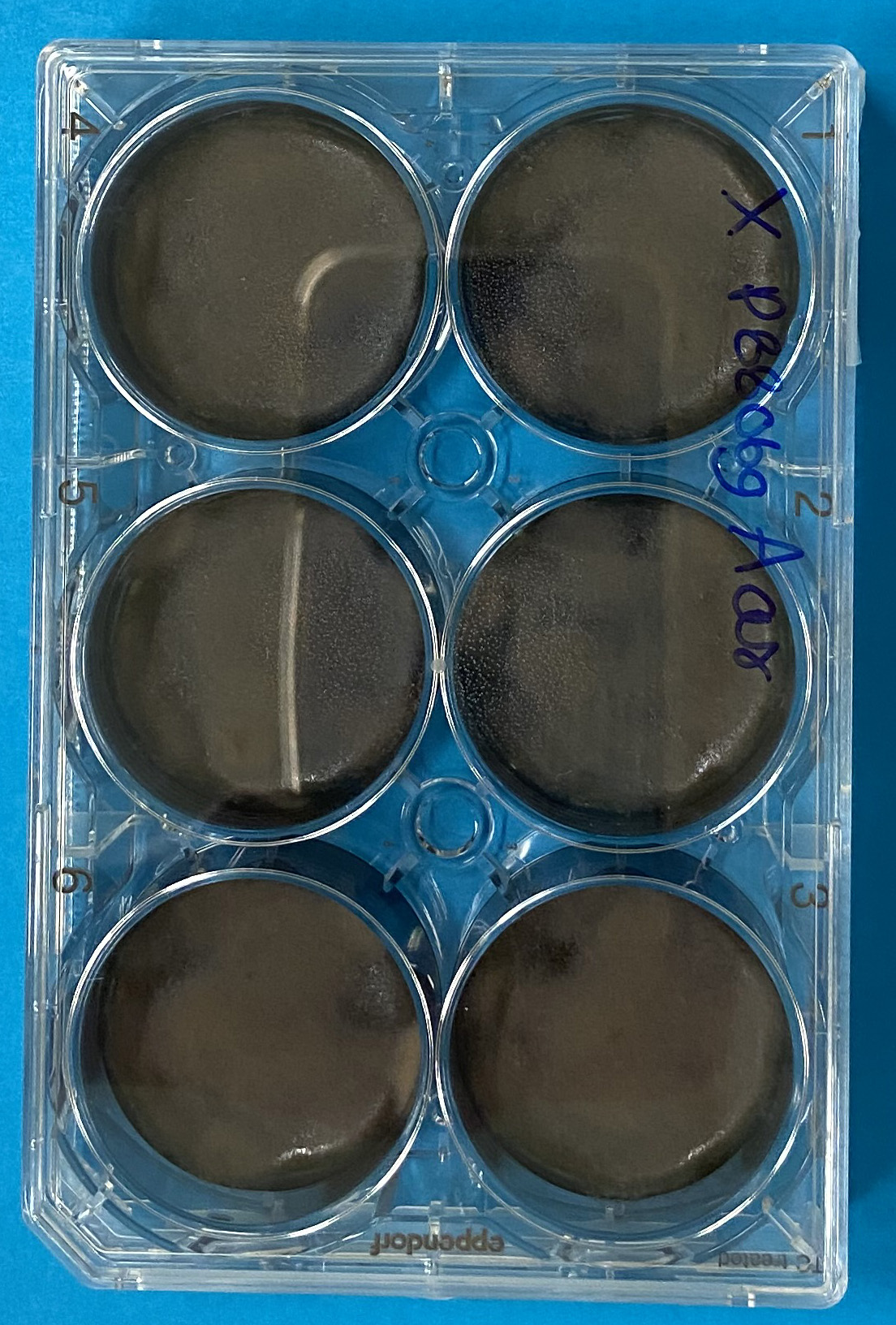

### Supplemental Figure 4

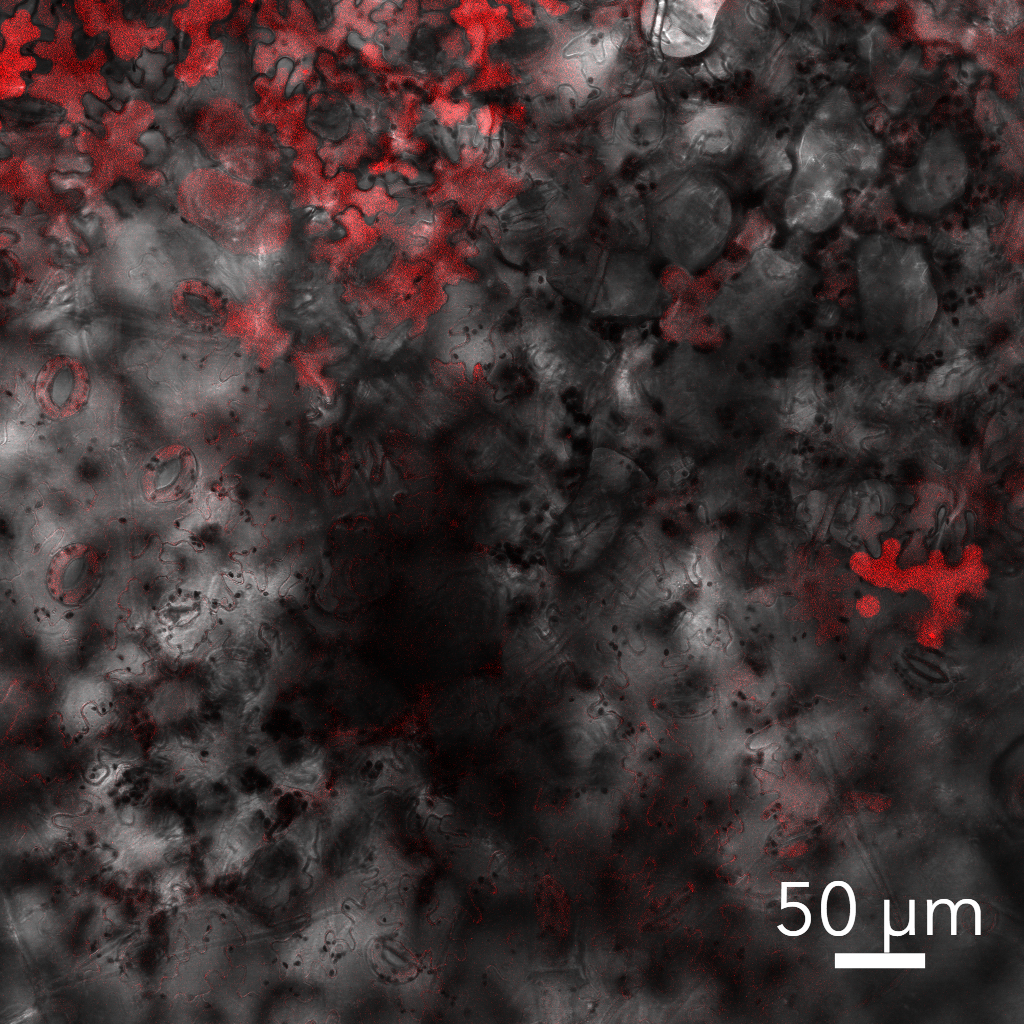
